# Correlative Organelle Microscopy: fluorescence guided volume electron microscopy of intracellular processes

**DOI:** 10.1101/2021.03.29.437317

**Authors:** Sergey Loginov, Job Fermie, Jantina Fokkema, Alexandra V. Agronskaia, Cilia de Heus, Gerhard A. Blab, Judith Klumperman, Hans C. Gerritsen, Nalan Liv

## Abstract

Intracellular processes depend on a strict spatial and temporal organization of proteins and organelles. Directly linking molecular to nanoscale ultrastructural information is therefore crucial to understand cellular physiology. Volume or 3-dimensional (3D) correlative light and electron microscopy (volume-CLEM) holds unique potential to explore cellular physiology at high-resolution ultrastructural detail across cell volumes. Application of volume-CLEM is however hampered by limitations in throughput and 3D correlation efficiency. Addressing these limitations, we here describe a novel pipeline for volume-CLEM that provides high-precision (<100nm) registration between 3D fluorescence microscopy (FM) and 3D electron microscopy (EM) data sets with significantly increased throughput. Using multi-modal fiducial nanoparticles that remain fluorescent in epoxy resins and a 3D confocal fluorescence microscope integrated in a Focused Ion Beam Scanning Electron Microscope (FIB.SEM), our approach uses FM to target extremely small volumes of even single organelles for imaging in volume-EM, and obviates the need for post correlation of big 3D datasets. We extend our targeted volume-CLEM approach to include live-cell imaging, adding information on the motility of intracellular membranes selected for volume-CLEM. We demonstrate the power of our approach by targeted imaging of rare and transient contact sites between endoplasmic reticulum (ER) and lysosomes within hours rather than days. Our data suggest that extensive ER-lysosome and mitochondria-lysosome interactions restrict lysosome motility, highlighting the unique capabilities of our integrated CLEM pipeline for linking molecular dynamic data to high-resolution ultrastructural detail in 3D.

**Significance:** We have developed a correlative imaging pipeline to **(i)** correlate 3D-FM to volume-EM data with high precision, directly bridging the FM and EM resolutions **(ii)** achieve high-throughput volume-CLEM by targeted EM imaging of a single organelle sized region-of-interest, pre-identified by FM **(iii)** link live-cell fluorescence imaging of cultured mammalian cells to high-throughput volume-CLEM **(iv)** quantitatively study structure-function relations at subcellular scale **(v)** link rare (e.g. membrane contact sites) and transient (e.g. organelle interactions) cellular events to 3D ultrastructure.

The targeted volume-CLEM pipeline provides a unique prospect for multi-modal correlative intracellular analysis combining dynamic interaction (live-cell imaging), functional state (live-cell imaging), molecular localization (FM), and 3D-ultrastructure (FIB.SEM) at nanometer scale.

## Introduction

Eukaryotic cells are compartmentalized in organelles delimited by distinctive intracellular membranes, each with special biochemical functions, yet functioning together to maintain cellular homeostasis. Complications in organelle performance are associated with many pathologies ranging from infection, neurodegeneration, and cancer (Ballabio & Bonifacino, 2019; Ferguson, 2018b; Sironi et al., 2020; Wu et al., 2018). A next step in understanding cellular regulation is how interconnectivity between different types of organelles is important for function (Gatta & Levine, 2016). This calls for novel microscopy approaches to study spatial and temporal regulation of intracellular processes at the nanoscale level.

Electron microscopy (EM) has immensely contributed to nanoscale understanding of complex intracellular structure that underlies diverse cellular functions. The biggest challenge for EM, however, has been the characteristic lack of 3D information, arising from the limited section thickness obtained by classical transmission electron microscopy (TEM). Building on the initial idea of collecting serial TEM or electron tomography (ET) images of consecutive sections to build a 3D volumetric reconstruction (Keene et al., 2008; Knott et al., 2008; Saalfeld et al., 2010), various volume-EM approaches have now been developed. Scanning EM (SEM) based volumetric approaches; array tomography, in which serial sections from the sample are collected on a substrate (Micheva & Smith, 2007), serial blockface SEM (SBF.SEM), in which the blockface is imaged repeatedly after sectioning by an in-situ ultramicrotome (Denk & Horstmann, 2004), and focused ion beam SEM (FIB.SEM), where an ion beam removes slices from the blockface (Heymann et al., 2006). Especially FIB.SEM, which offers isotropic sub-5nm resolution in x, y, and z, allows to gather 3D EM information at resolutions able to address many significant biological questions (Bushby et al., 2011; Kizilyaprak et al., 2014; Knott et al., 2011; Lucas et al., 2012; Narayan et al., 2014; Christopher J Peddie & Collinson, 2014). Volume-EM data also opens up the possibility of quantitative EM studies without the dimension restriction of 2D EM, by automizing collection of 3D ultrastructural data. Thus, the stage is ready for volume-EM to deliver its promise in cell biology (Hoffman et al., 2020), just like confocal fluorescence microscopy (FM) did in the last decades by enhancing our understanding of 3D organization of molecules and organelles within cells.

The unique potential of volume-EM is currently limited in throughput as for each experiment a relatively large cellular volume is imaged at high-resolution. This is incompatible with most life sciences studies, which often require analysis of small volumes of several independent samples. If the high-resolution acquisition could be limited to a pre-defined region of interest (ROI), throughput of volume-EM would be considerably enhanced (Burel et al., 2018; Ronchi et al., 2021). With the introduction of correlative light electron microscopy (CLEM), a ROI can be highlighted/selected with FM (a common driver for CLEM). Volume correlative light and electron microscopy (volume-CLEM) approaches offer a unique potential to explore molecular characteristics together with high-resolution ultrastructural details across the cell volume. Recent studies, especially on connectomics, provide very promising examples of volume-CLEM for tissue analysis (Collman et al., 2015; Oberti et al., 2011), for example by visualizing glia in 3D reconstructions of mouse hippocampal tissue at nanometer resolution (Fang et al., 2018). Also in cellular level, state-of-art volume-CLEM studies visualize protein-ultrastructure relationships in three dimensions across whole cells by identification of morphologically complex structures within the crowded intracellular environment (Hoffman et al., 2020). However, an accurate and reliable correlation between separate 3D-FM and 3D-EM platforms data sets is far from trivial because of the resolution mismatch and the sample transfer in between the two modalities (Ronchi et al., 2021). Current efforts in volume-CLEM development focus around improving its accuracy, throughput and accessibility. An integrated CLEM platform, i.e. combining FM and EM in one instrument, inherently resolves this correlation problem since the coordinate planes of the FM-EM are shared. Recently introduced integrated confocal-FM and volume-EM systems, which have the capability to record both confocal fluorescence and reflection images (Ando et al., 2018; Brama et al., 2016), can be used to achieve a robust and streamlined image acquisition. 3D-FM can identify 3D coordinates of ROIs within cells, which can be traced and imaged in volume-EM in a targeted way, directly increasing the throughput of the method. Correlation precise enough to target a sub-cellular level ROI (e.g. an organelle) in 3D is specifically challenging and has thus far not been demonstrated.

Understanding intracellular processes e.g. organelle interconnectivity ultimately requires linking molecular and dynamic information from live-cells to high-resolution ultrastructure. Therefore, a next major step forward in volume-EM will be to correlate EM data to functional information from (live-cell) FM (Blazques-Llorca et al., 2015; Hoffman et al., 2020). We and others have shown that linking functional or dynamic information obtained with live-cell imaging to the underlying fine structure of the cell opens up powerful possibilities to study mechanistic processes with respect to their ultrastructure (Collinson et al., 2017; Fermie et al., 2018; Russell et al., 2016). Correlation with live-cell FM also aids the identification and capture of rare cellular structures or events, which is very challenging, time demanding, and at times basically impossible without smart tracking for EM imaging (Burel et al., 2018; Delpiano et al., 2018). An integrated CLEM platform provides exactly these requirements by enabling a direct translation between live-cell FM and volume-EM and the targeting identified live-cell events for follow-up ultrastructural imaging. Therefore, an FM-guided imaging pipeline for volume-EM data collection is essential to improve the efficiency, throughput, and quantitative capabilities of the technique.

Here, we present an optimized imaging pipeline to identify and target single organelles for volume-CLEM within the complex cellular environment. Following live-cell FM and fixation, we prepare cells for volume-EM imaging. We then use confocal laser scanning microscopy integrated in a FIB.SEM to quickly and accurately re-trace single endo-lysosomes in the complete sample, and generate the corresponding 3D volume-EM data set. With this pipeline we correlate motile characteristics of targeted organelles to their morphology, yielding essential information on their identity and cellular surroundings, such as number and types of contact sites with the endoplasmic reticulum (ER).

## Results

### Integrated 3D CSLM / FIB.SEM pipeline for deep sub-cellular precision volume-CLEM

Correlating single organelle sized, intracellular regions-of-interest (ROIs) from fluorescence (FM) to electron microscopy (EM) is challenging due to the limited resolution of FM, the inherently different contrast mechanisms of the two modalities and highly crowded content of the cell (Ando et al., 2018). These challenges are amplified when correlating 3D FM to 3D EM images, and even further when using live-cell imaging as FM method. Integrated CLEM instruments (with the LM built in the EM) greatly facilitate retracing of the ROI from FM to EM (Koning et al., 2019; Liv et al., 2013), since the coordinate systems of the FM-EM are shared. Recently, multiple integrated (confocal) FM and volume-EM systems were reported (Ando et al., 2018; Brama et al., 2016; Delpiano et al., 2018; Gorelick et al., 2019; Lane et al., 2019). We add to this a home-built system integrating a Confocal Laser Scanning Microscope (CLSM) into a Focused Ion Beam / Scanning Electron Microscope (FIB.SEM), similar in geometry to a previously reported (Timmermans et al., 2016) integrated system and as outlined in Supplementary Figure 1. In short, the CLSM unit is mounted on one of the side ports of the FIB.SEM chamber. The FIB.SEM and CLSM image the sample from the same direction, while switching between the two modalities is simply accomplished by shuttling the specimen on the accurate motorized stage. The integrated CLSM (iCLSM) is equipped with a Nikon industrial inspection objective lens (ELWD series, Plan Apo 100x NA0.9).

### Endocytic fiducials for improved 3D correlation

Even in such an integrated CLEM platform, the accuracy of FM-EM registration/ correlation is limited by FM resolution, especially in the z-dimension. Several organelles can be located within the same fluorescent spot, causing the risk of misidentification. We tackled this problem by adding gold core silica shell fiducials to cells prior to imaging. These fiducials, recently developed in our labs (Fokkema et al., 2018; Mohammadian et al., 2019), are visible and highly compatible with live-cell imaging. They are taken up by endocytosis, which generates a natural 3D distribution within the cell, throughout the endo-lysosomal system (Fokkema et al., 2018; Prabhakar et al., 2020). The resulting array of well-distributed puncta provides an accurate 3D translation map between FM and EM, and enables identification and registration of even single organelles. Moreover, their fluorescence is retained after osmium fixation, and serve as suitable anchors for the translation of coordinate systems between stand-alone CLSM, iCLSM, and volume-EM.

We then set up the following high accuracy, live-cell volume-CLEM pipeline: image live cells in a stand-alone FM – fix cells and make a confocal Z-stack – prepare cells for FIB.SEM – place and image the sample in the integrated platform-map confocal Z-stack to 3D iCLSM image - target and image the pre-identified region in FIB.SEM. A detailed outline of this high accuracy live-cell volume-CLEM workflow is shown in **Figure 1**.

**Figure 1.**
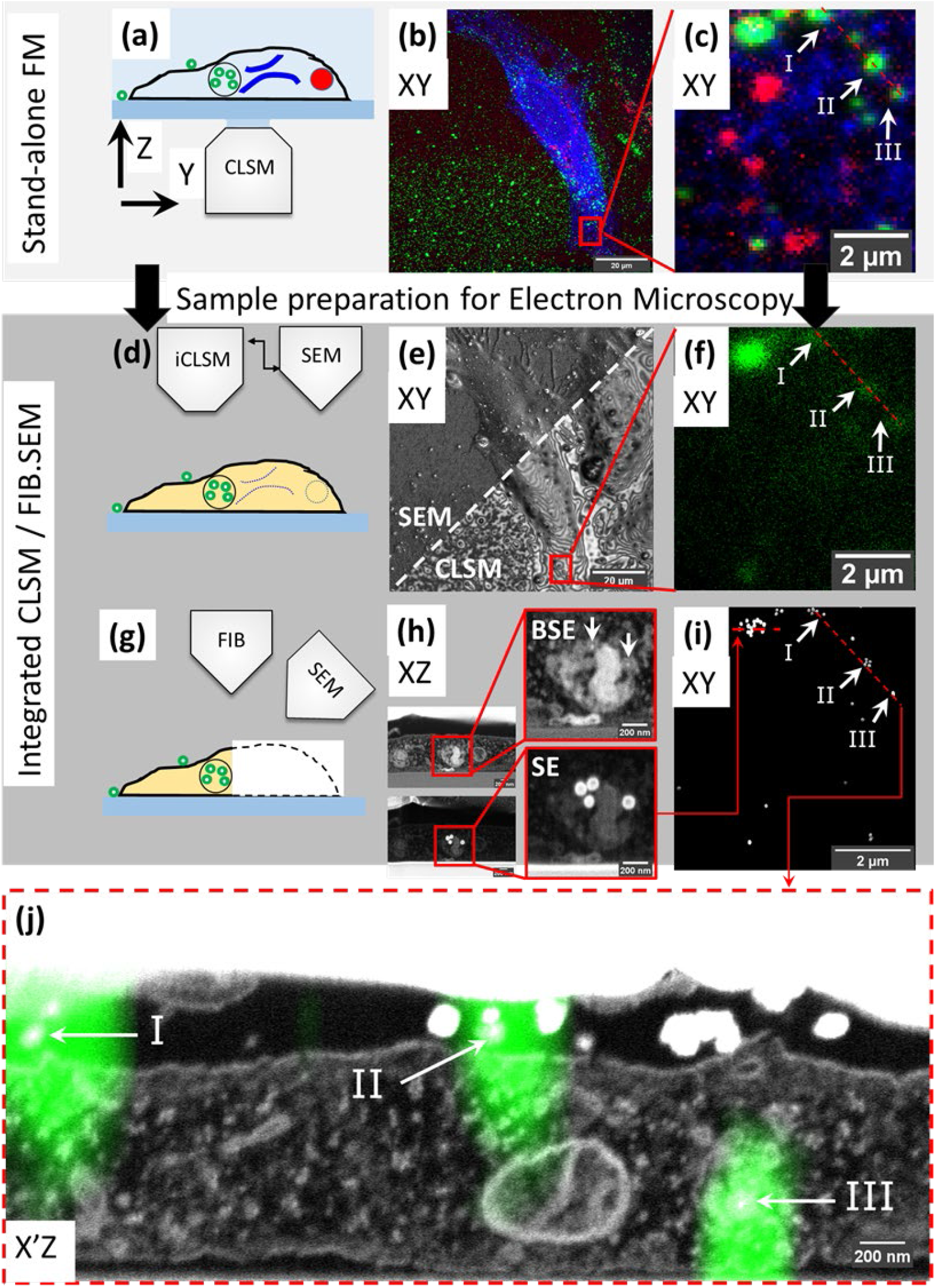
High accuracy 3D correlation workflow from live-cell-to-volume-EM with iCLSM. **(a-c) Live-cell imaging and CLSM. (a)** Live cell imaging of cells on gridded coverslips in stand-alone CLSM followed by z-stack recording of fixed cells. **(b)** Maximum intensity projection of the CLSM Z-stack. ER (blue), active lysosomes (red), and endocytic fiducial particles (green). **(c**) the selected ROI, red square, within the cell in (b). **(d-f) Correlation with iCLSM. (d)** Locating the ROI in the FIB.SEM chamber by iCLSM after EM preparation. **(e)** Reflection mode imaging in iCLSM to navigate through the gridded glass coverslip, and fast tracking of ROI by cell shape. iCLSM reflection image (lower half) relates easily with SEM (upper half). **(f)** The ROI can be localized by the fluorescence mode of iCLSM. The (remaining) fluorescence signal of the fiducial nano-particles is used for correlation back to the CLSM image in (c). **(g-i) 3D FIB.SEM acquisition of the re-established ROI. (h)** Slice-by-slice acquisition is performed with backscattered-electrons (BSE, top) and secondary electrons (SE, bottom). Fiducial nano-particles have a clear signature of a bright backscattering core (top) and shell in secondary electrons (bottom). **(i)** They can be thus filtered from the acquired data stack to aid precise correlation of ROI between CLSM and FIB.SEM. Note the one-to-one correspondence between panels (c)-(f)-(i). **(j)** Overlay of 3D CLSM data from the fixed cell and the 3D FIB/SEM data. EM image corresponds to the plane of red dashed line in (c)-(f)-(i).

### Fluorescent microscopy; from live-cells to 3D confocal Z-stacks

To demonstrate our approach we labeled live HeLa cells for 2 distinct organelles; Endoplasmic Reticulum (ER) and lysosomes. Cells were cultured on gridded glass coverslips with etched marks (Polishchuk et al., 2000) and transiently transfected (16 hours) with mEmerald-Sec61β, localizing to ER membranes. Then, the transfection medium was replaced with medium containing SiR-lysosome (SiRLyso) (Lukinavičius et al., 2016), marking functionally active lysosomes, and containing the above introduced endocytic fiducial nanoparticles, both for 3 hours at 37°C. To view the dynamics of ER and lysosomes and their interactions we performed live-cell imaging of individual fluorescent organelles. We recorded the mEmerald (ex/em 487/509nm) and the SiRLyso (ex/em 652/674nm) channels with 1s/frame over 2-3 minutes (Figure 1a). Then cells were fixed *in situ* on the microscope stage by adding fixative directly in the imaging chamber. This assures that there is no imaging gap between the last live-cell frame and the corresponding image after fixation. Following fixation, fluorescent confocal Z-stacks were recorded of the same cell capturing the 3D distribution of mEmerald-Sec61β, SiRLyso and the fiducial particles (ex/em 543/576nm). Figure 1b shows a maximum intensity projection of the Z-stack; fiducial particles (green), ER (blue) and lysosomes (red) are visible. With the confocal Z-stack, the exact x-y-z coordinates of ER and lysosomes are visualized with respect to the fiducial markers (Figure 1c).

### Relocating the ROI in FIB.SEM using iCLSM

Samples were removed from the microscope stage, post-fixed, with osmium-thiocarbohydrazide-osmium (R-OTO), and further stained with uranyl acetate and Walton’s lead aspartate. Cells were embedded in resin following the ‘extremely thin layer plastification’ (ETLP) method (van Donselaar et al., 2018). For a more in depth description of these protocols we refer to the Methods section. After resin embedding, the cells were coated with carbon, mounted on a stub, and transferred to the integrated CLSM/FIB.SEM set-up (see Supplementary Figure 1). A quick re-localization of the cells imaged previously in the stand-alone CLSM was achieved by using the SEM to show the markings on the coverslips (Figure 1d-f). The confocal reflection image of these same marks using the iCLSM allowed easy correlation between the CSLM to find back the cells (Figure 1e). The reflection light images expose various surface topology features of the cells. As many of these features are also visible in the SEM images, this registration generates a first, 2D, correlation map around the ROI (red rectangle in Figure 1b and 1e). Then the confocal fluorescence image from iCLSM localizes fiducial nanoparticles (which notably retain their fluorescence in the epoxy resin) in x, y, and z (Figure 1f). iCLSM image of the fiducial particles directly matches the fiducial channel in the CLSM image collected before EM sample preparation (note the correspondence between Figure 1c and 1f), and provides a 3D translation anchor. After this, we could reliably start the 3D FIB.SEM acquisition of this targeted ROI.

### Volume-EM imaging and 3D correlation

In FIB.SEM, samples are imaged by scanning the surface of a ROI using the electron beam, after which a thin layer is ablated from the surface using the FIB (Figure 1g). This cycle is repeated until the ROI has been imaged, allowing a 3D reconstruction of the sample. We performed the slice-by-slice acquisition both with backscattered-electrons (BSE) (Figure 1h, top) and secondary electrons (SE) (Figure 1h, bottom). Fiducial nanoparticles have a clear signature, with a bright backscattering by both the gold core and a bright secondary electron emission by the silica shell, which improves their visibility and makes them well-suited for FIB.SEM image acquisition. The fiducial nanoparticles are found in endo-lysosomal compartments, and can be detected at an individual particle level (Figure 1h). With the high resolution provided by FIB.SEM in x, y and z dimensions, single particles can be identified, and fitted to the fluorescence Z-stack data (Figure 1j). Using the fiducials we correlated and registered the 3D CLSM data from the fixed cell with the 3D FIB.SEM data set (Figure 1i and j, also see Supplementary 3D-overlay Video 1). Figure 1j shows the reconstructed EM image corresponding to the dashed line in the FM images in Figure 1c, and f. The fluorescent spots I and II shown with arrows point to fiducials outside the cell and III to intracellular fiducials in an endosome. We show a correlation accuracy at the level of single, 90 nm-sized nanoparticles, which is far below the x, y, z resolution limits of the FM, and allows to study sub-organelle structures.

### Correlation of organelle motility and function to morphology by targeted volume-CLEM

Next we utilized our high-precision, live cell to volume-CLEM pipeline to link dynamic behavior of single, active endo-lysosomes to their morphology and neighboring environment at nanometer resolution. Besides motility, live-cell imaging was used to analyze functional characteristics of organelles, exploiting fluorescent reporter probes. As an example we used the SiRLyso probe to report the activity state of lysosomal hydrolase Cathepsin D in single lysosomes (Butkevich et al., 2017). In a previous 3D CLEM study (Fermie et al., 2018), we used a constellation pattern of at least 3 fluorescent spots found by live-cell imaging, to retrace a similar pattern of endosomal organelles in the volume-EM data. Although this proved a powerful approach, correlation of 3D FM and 3D EM data by matching the organelle constellations was relatively time consuming and limited in accuracy. With our current approach, correlation of the fiducial signal between the pre-embedding CLSM z-stack and post-embedding iCLSM assures that we know the exact z-plane of each organelle imaged in live-cell FM. Moreover, our correlation is not limited to organelles bearing fiducial particles, but extends to all fluorescent signals, even when lost after EM embedding, using the fluorescence signal from fiducial particles as an anchor in translating data to volume-EM. Hence, we can correlate each live-cell imaged organelle to the volume-EM data. This significantly eliminates the time and computational need for correlation of matching organelle constellations between datasets, and hence notably increases the throughput (e.g. organelles analyzed per cell) of the live-cell CLEM workflow.

Live-cell imaging allows to study key temporal, functional (e.g pH and hydrolase activity as we show here) and structural parameters of single organelles over an extended period of time, and by 3D CLEM we correlate these directly to nanometer architecture, cellular context, and inter-organelle connections. As a proof of principle **Figure 2** shows the correlation sequence from a live-cell movie of hydrolase active lysosomes and ER to volume-EM. For live-cell imaging (Figure 2a), we recorded SiRLyso (lysosomes) and Sec61β (ER) channels in a single focal plane to reach the temporal resolution required to visualize transient events on a sub-second scale (<1s between frames, see Supplementary live-cell Video 2). We analyzed 14 lysosomes (SiRLyso positive organelles) for their dynamic behaviors, such as speed, displacement, fusion or interaction with other lysosomes and ER (Figure 2b). We then fixed cells *in situ* by adding fixative directly to the medium in the live-cell holder, while the camera was still acquiring images (Figure 2c). In the fixed material, Z-stacks of the ROI were recorded to visualize SiRLyso, Sec61β and fiducial particles (Figure 2c). The samples were prepared for EM, and imaged following the routine explained in the previous section and Figure 1. With the ultrastructural resolution of EM, we analyzed compartment identity, fusion profiles with other compartments, interactions with surrounding structures, and inter-organelle interactions. Finally, we integrated the multi-modal data collected per organelle, from live-cell imaging to 3D-EM (see Supplementary 3D-overlay Video 1).

**Figure 2.**
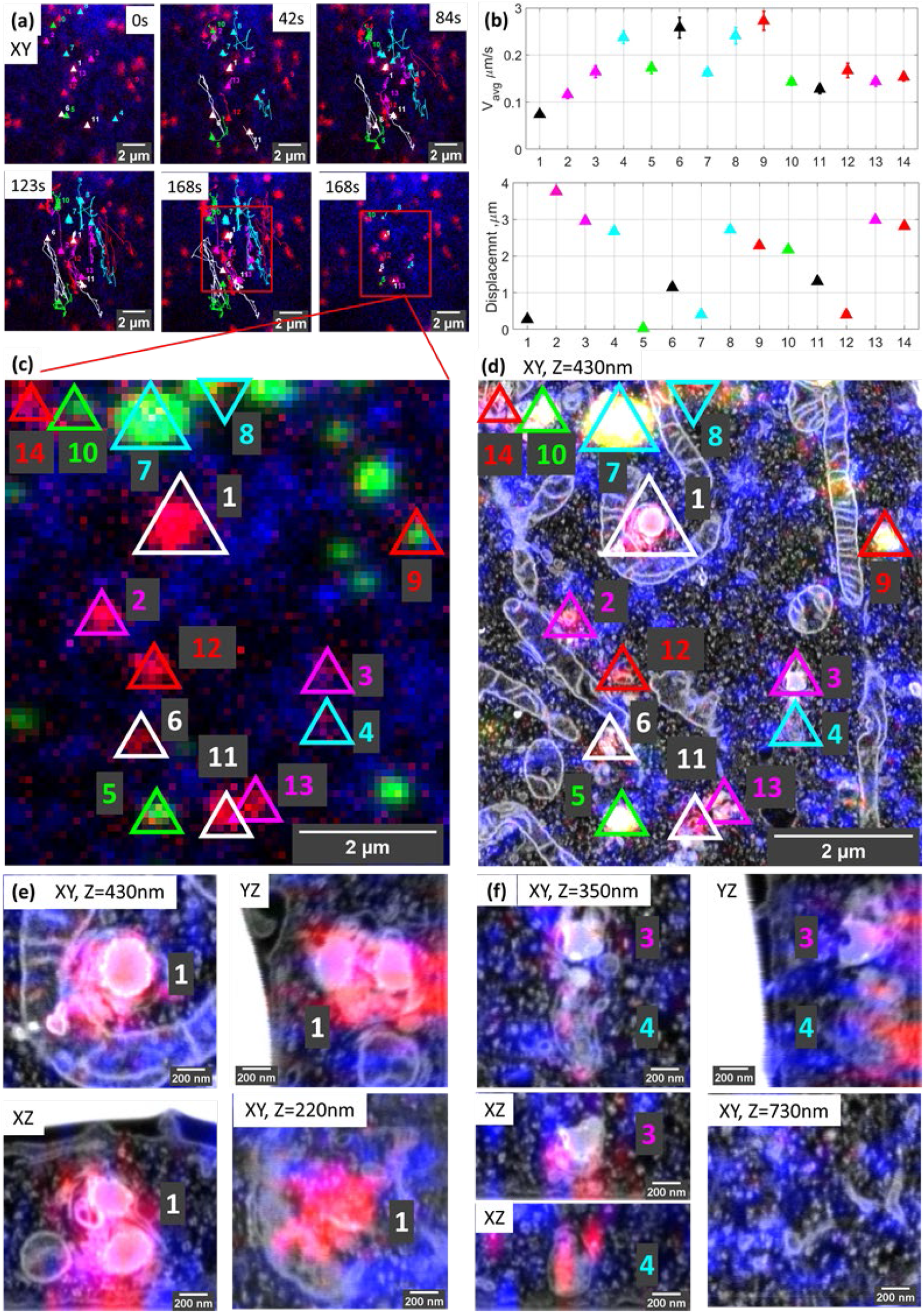
Organelle dynamics directly correlated to morphology. **(a)** Time-lapse images from live-cell imaging. Lysosomes are stained with SirLyso (red), ER with mEmerald-Sec61β (blue) and the cells have endocytosed fiducial particles (green). We selected 14 SirLyso positive - enzymatically active lysosomes and traced their dynamic behaviour over 3 minutes. **(b)** Plot of the average velocity (µm/s) and total displacement of each selected organelle, numbered from lysosome#1 to lysosome#14 in the x-axis. For example, lysosome#1 is very static, whereas lysosome#4 moves fast over extended distances. (**c)** Max intensity projection of CLSM stack recorded on the same cell after fixation. Fiducial markers are visible in green. Each identified and analysed organelle is numbered as in (b). **(d)** Following live-cell imaging and CLSM, cells are prepared for EM, imaged, and high precision correlation is achieved using iCLSM-FIB.SEM. **(e)**. Higher magnification images of lysosome#1, lysosome#3, and lysosome#4 display their distinct morphologies from different planes and interaction with surrounding cellular organelles. See supplementary data for the rest of the lysosomes.

The movies defined lysosome#1 as the most stable organelle of the whole set; it shows no displacement within 3 minutes and its velocity is fixed at 0.07 µm /s, resembling Brownian motion in a confined area. Lysosomes are heterologous in shape and by EM can be classified by morphological characteristics such as size and shape, as well as electron density and contours of their content (Fermie et al., 2018; Meel & Klumperman, 2008). The EM data identified Lysosome#1 as an (auto)lysosome, showing a heterogeneous content with an electron dense region and irregularly organized internal membranes. Its size is 0.71×0.69×0.74 µm. Intriguingly, this stable lysosome has extensive contact sites, covering two-thirds of its surface area, both with mitochondria and ER. Especially its interaction with the mitochondria wrapping it from 3 sides over a 2 µm^2^ contact area creates a very confined space (Figure 2d-e).

In contrast to Lysosome#1, Lysosome#4 was very motile with an average velocity of 0.2 µm /s, traveling over a distance of 2.8 µm in 3 minutes before it reached the final position visualized also in EM. The organelle first exhibits directional movement towards (∼100°) the nucleus, then meets lysosome#3 (which is quite stable until the meeting point) at 150s, after which they together move directionally towards (∼270°) the plasma membrane (see Supplementary live-cell Video 2). The directional and stable trafficking characteristics indicate that lysosome#3 and lysosome#4 are transported via microtubules and associated motors. The size of lysosome#3 is 0.43×0.61×0.51 µm, and by EM it shows a heterogeneous dense lumen. Lysosome #4, with a size of 0.39×0.77×0.54 µm, on the other hand, has a relatively electron-lucent lumen compared to lyososomes#1 and #3, with irregularly organized internal membranes and numerous intraluminal vesicles (ILVs) (Figure 2d-f). In contrast of lysosome#1, neither lysosome#3 nor lysosome#4 display contact sites with mitochondria and are not wrapped by ER cisternae. Rather, they only touch the ER at 2 and 3 points, respectively. As also reported by others (Cabukusta & Neefjes, 2018; Wong et al., 2019), these data indicate that interactions of lysosomes with other organelles have a defining role in their motile characteristics. Our high accuracy live-cell volume-CLEM workflow provides a unique means to mechanistically study these interactions. For further analysis of the organelles in this ROI see Supplementary Figure 2. Note that 10 out of the 14 organelles correlated and analyzed in the ROI have no fiducial particles.

### Rapid volume-CLEM of targeted subcellular ROIs with <100 nm accuracy

The possibility of matching all coordinate planes between 3D-FM and 3D-EM modalities allows us to address a major bottleneck in volume-EM approaches: the throughput. In most volume-EM imaging techniques, especially in FIB.SEM, hitting the ROI within the imaging volume is ensured by maximizing the image area. This imposes a compromise between long imaging durations (days to weeks) and resolution (low to high). Here, we tackle this problem by using iCLSM to reliably identify and select a very confined ROI (e.g. a single lysosome) for volume-EM data collection. Using this targeted approach, we lower the imaging time for a selected ROI from several days to 1-2 hours.

To demonstrate the power of the targeted imaging approach in **Figure 3**, we selected a very small ROI of a single SiRlyso positive lysosome devoid of fiducial particles (Figure 3a and b, red square) for volume-EM. Using the endocytic fiducials in the surrounding organelles as anchor, the CLSM - iCLSM correlation provides an accurate 3D map between FM and EM and delivers the 3D coordinates of all organelles within the complete block, including the selected organelle without fiducial particles (e.g. the red labelled lysosomes in Figure 3a are not visible in Figure 3b). As the x, y, and z coordinates of the identified organelles are exactly known a priori, only a very small volume needs to be imaged in FIB.SEM. This greatly minimizes the time spent on pre-imaging procedures (i.e. trench milling and Pt layer deposition, Figure 3c), as well as the actual image acquisition. Hence, the targeted lysosome was visualized in 3D within only ∼2.5 hours in FIB.SEM (with voxels of 1.2×1.2×10 nm and dwell time of 3 µs), by direct correlation to the corresponding FM data (Figure 3d, see also Supplementary 3D-overlay Video 3). The CLEM data show by FM that the selected organelle is positive for active Cathepsin D (SiRLyso signal) and not reached by endocytosed fiducial particles. By EM, we confirm the absence of fiducial particles and show that the lysosome has an electron dense lumen with clearly degraded content, a spherical shape with an approximate diameter of 560 nm, and extensive contact sites with ER and mitochondria.

**Figure 3.**
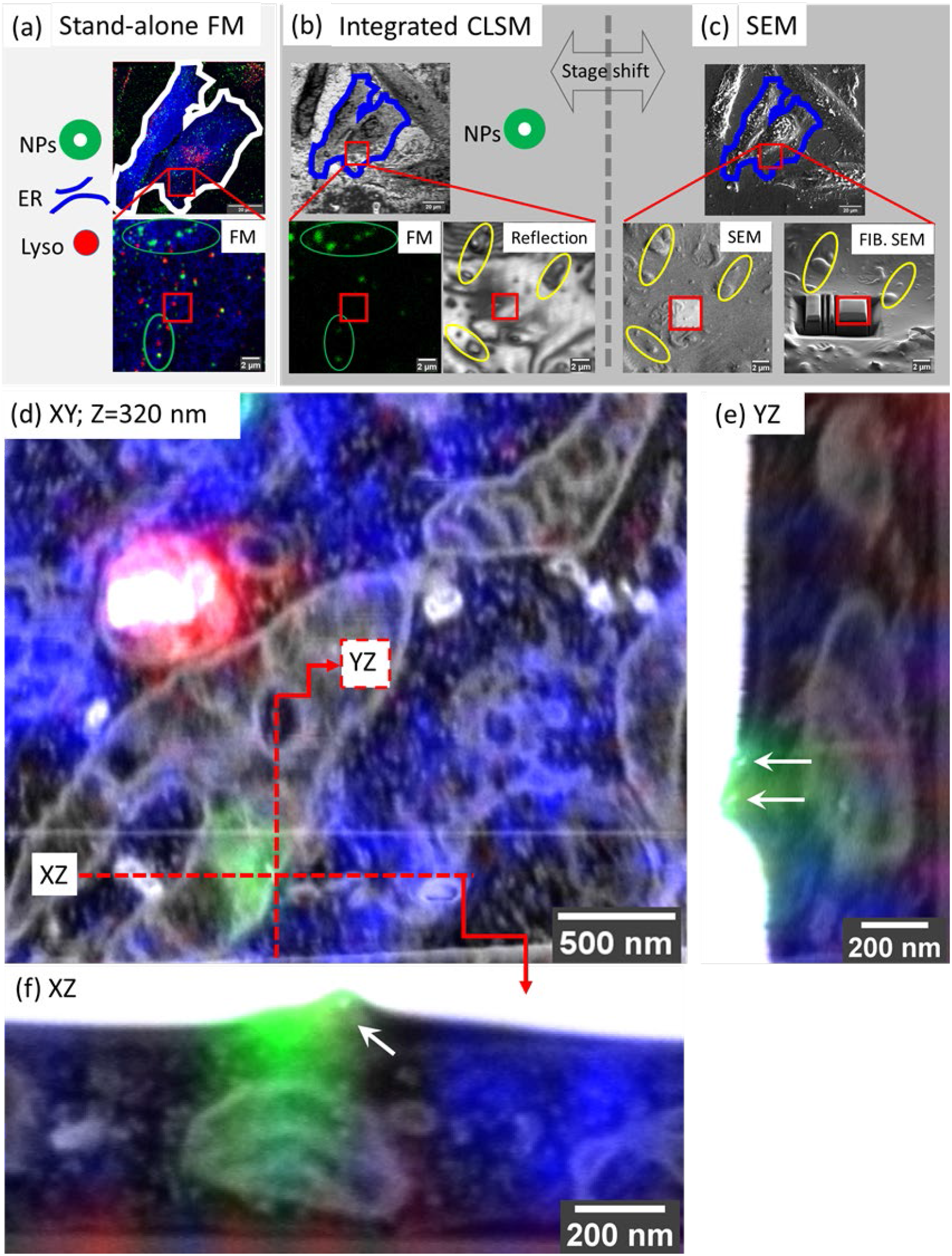
Targeted imaging of minimized volumes in FIB.SEM using iCLSM. (a-c) High accuracy correlation routine for targeted imaging in FIB.SEM. **(a)** Maximum intensity projection of a confocal z-stack collected at CLSM from fixed cells, following optional live-cell imaging. 3D CLSM data shows ER (mEmerald-Sec61β, blue), lysosomes (SirLyso, red), and endocytic fiducials (green). Red square indicates the ROI bearing a single lysosome selected for targeted volume-EM. **(b)** The cells and the ROI are traced back after EM sample preparation. Fluorescence of endocytic fiducials is visualized by iCLSM and correlated to the CLSM data (pink ellipses), providing the coordinates of the ROI, even when the ROI itself has no fiducial particles. Reflection image collected by iCLSM provides topographic information which is correlated with SEM for fine alignment (green ellipses). **(c)** Once precisely located, the Pt layer is deposited to protect the ROI from FIB, and trenches are prepared for imaging only the small ROI area. **(d)** The small ROI, bearing a single lysosome not reached by endocytic fiducial particles, is targeted and imaged in FIB.SEM. 3D CLSM data, ER, lysosomes, and endocytic fiducials, is overlaid with the 3D FIB.SEM data. **(e, f)** 2 fiducial particles located on the plasma membrane are used as a blind target to assess correlation/targeting efficiency. Green fluorescence signal of the particles is correlated to their gold core with an accuracy of 60 nm in X **(e)**, 90 nm in Y **(f)** and 330 nm in Z.

To measure our correlation precision with high accuracy, we next used a single fluorescent spot of the fiducial particles as a blind target. The faint green fluorescence signal visible in Figure 3d, originating from another z-plane (orthogonal planes depicted with the red lines), belongs to fiducial particles attached to the cell membrane (orthogonal views shown in Figure 3e,f). Blind correlation of the signal recorded from this single spot in CSLM, i.e. before EM preparation, with the volume-EM data, placed the peak of the fluorescence signal 60 nm in X, 90 nm in Y and 330 nm in Z direction from the centre of mass of the two fiducial particles. This indicates that we can reach a correlation accuracy of sub 100 nm in the XY dimension and sub-350 nm in Z direction.

Together these data show that our high precision volume-CLEM approach allows to target single organelles identified CLSM, which within 1-2 hours can be correlated, targeted and imaged in volume-EM with 100 nm confidence. This reliable targeted imaging approach, greatly improves the throughput of volume-EM technique.

### Targeted volume-CLEM efficiently reveals inter-organelle contact sites

The possibility of rapid, targeted volume-CLEM with < 100 nm precision opens new avenues for imaging at sub-organelle scale (e.g. organelle subdomains), as well as the correlation of rare or transient structures in cells. As an illustration for this application, we examined membrane contact sites (MSCs) between lysosomes and ER. Organelles can communicate with each other by vesicular traffic as well as by MCSs, by which membranes are closely positioned and tethered, but there is no fusion. This special case of intracellular communication facilitates metabolic channeling between distinct (not homotypic) organelles. MSCs between ER and lysosomes mediate the exchange of signaling molecules, ions, metabolites and lipids, which is important for endo-lysosome maturation and positioning (Friedman et al., 2013; Hönscher et al., 2014; M. L. L. M. Jongsma et al., 2016; Wijdeven et al., 2016). Only recently, the importance of MScs in cell physiology was fully recognized, which has made them a focus of attention in contemporary cell biology studies (Phillips & Voeltz, 2015; Wu et al., 2018). However, studying MSCs with high spatial and temporal resolution remains a challenge because of their small size and transient and confined nature. ER-lysosome MSCs are identified as closely apposed (<20 nm) membranes over a distance of 20nm (Huang et al., 2020; Scorrano et al., 2019). Hence, (3D) EM is essential to provide sufficient resolution to examine the presence and structure of these inter-organelle contacts. However finding MSCs by EM is akin to seeking a needle in the hay stack. Here we employ our targeted imaging pipeline to efficiently identify MSCs between ER and lysosomes in live-cells and visualize their corresponding ultrastructure in 3D (**Figure 4**).

**Figure 4.**
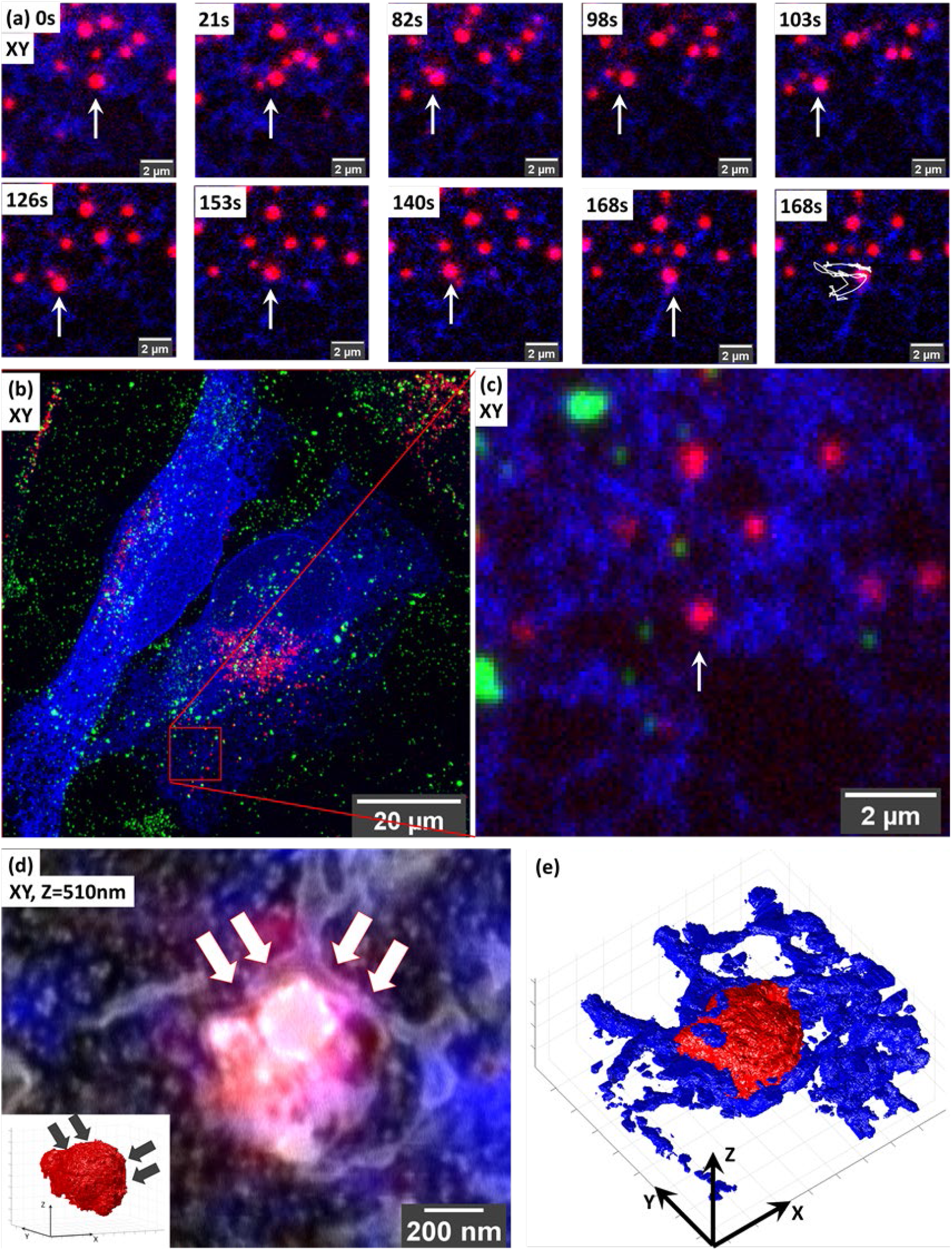
Targeted volume-CLEM of ER-lysosome contact sites implicates ER in lysosomal motility. (a) Live-cell imaging of lysosome dynamics in relation to ER contacts. Lysosomes are stained with SirLyso (red), ER with mEmerald-Sec61β (blue). Arrow points to the lysosome targeted for imaging. At 168s, the last image of the time-lapse panel, the dragon tail shows that the lysosome mainly follows the ER tubules, and that ER contact on its right side confines its movement. (b) Maximum intensity projection of the CLSM stack recorded after in-situ fixation, note the endocytic fiducial markers visible as green. (b) Image of entire cell indicating the imaged ROI (c) Zoom-in to the ROI. (d) Overlay of 3D CLSM data from the fixed cell and the 3D FIB.SEM data showing the single targeted lysosome traced back after resin embedding using iCLSM/FIB.SEM. The small ROI includes the lysosome-ER contacts. Inset shows that the curvature of the lysosome surface is altered on the site of dense ER interaction indicated with arrows. (e) Segmentation of the lysosomes (red), ER (blue) in the stack highlights the dense presence of ER on the right side of the lysosome.

We started this experiment in live cells in which we imaged a single lysosome (SirLyso) over 3 minutes for contact moments with ER (mEmerald-Sec61β) (**Figure 4**). In (live-cell) FM, even though the images are diffraction limited and it is not possible to make a direct conclusion, all lysosomes appear to have contact with ER. As shown in Figure 4a, the targeted lysosome exhibits a diffusive movement. First, it moves towards the left following the ER tubules (Figure 4a, 21s-98s), then it turns right but stops at a blockade formed by ER (Figure 4a, 98-103s). Then it turns left again and repeats this pattern (Supplementary live-cell Video 4). Figure 4b and 4c show the maximium intensity projection of the Z-stack. This analysis suggests that the dense ER visible on the right side of the lysosome defines a confined region for limited movement.

We then prepared the live-cell imaged cell for EM, and imaged the particular lysosome-ER interaction site following our targeted imaging pipeline. Note that this high-accuracy targeting is achieved without fiducial particles present in the targeted lysosome (Figure 4c). With the high confidence provided by our mehtod we could target a ROI of only **1.7** µm wide (Figure 4d) and collected 3D ultrastructural data in as little as 190 minutes between iCSLM and FIB.SEM (with voxels of 1.7×1.7×10 nm and a dwell time of 10 µs). Note the extremely small size of the imaged ROI compared to the CLSM field of view (132.45 µm wide) (Figure 4b, and c).

As presented in Figure 4d, the 3D-EM identified the targeted organelle as a lysosome with heterogeneous content, including electron dense material and irregularly organized internal membranes (Supplementary 3D-overlay Video 5). Its size is 0.66×0.69×0.50 µm. Importantly, the 3D-EM data showed that at its right side the targeted lysosome is fully covered with the ER cisternae, which follow the limiting membrane of the lysosome, conforming to its shape (Figure 4e, Supplementary 3D-segmentation Video 6 and Video 7). On its left side, we only found tips of ER cisterna touching the lysosome in a poking fashion. The ER on the left side showed a tubular thin lumen, whereas the right sided ER, which we know had a blocking effect on lysosome movement, exhibits a sheet-like thicker lumen (Figure 4d). Interestingly, the curvature of the lysosome on the right-side has a low convexity compared to its highly convex left-site (inset, Figure 4d), indicating the role of ER contacts also on the shape and possibly composition of lysosome membrane domains. These data show that ER forms multiples types contact sites with lysosomes, possibly with different functions, which in addition to exchange of biomaterials also defines lysosomal shape and movements. With this example we proof that our targeted CLEM approach provides a powerful tool to study MScs from live cells to 3D EM, with high temporal and spatial resolution, and a short processing time thanks to optimal ROI selection.

### Volume-CLEM of multiple interacting organelles followed over time

Lysosomes receive input from the endocytic, autophagic, phagocytic and trans-Golgi network pathways while they traffic throughout the cell (Saftig & Klumperman, 2009). They interact with other lysosomes and endosomes via kiss-and-run events or membrane fusion, resulting in the exchange of membranes and content (Huotari & Helenius, 2011; M. L. Jongsma et al., 2020; Pols et al., 2013). Although the essential features of lysosomal fusion events have been biochemically established, we don’t yet comprehend the spatial and temporal regulation of these processes (Ferguson, 2018a; Spang, 2016; van der Beek et al., 2019), simply because there is currently no means to analyze the ultrastructural background of multiple interacting lysosomes in an efficient way and in 3D. The targeted volume-CLEM pipeline tackles this challenge by allowing analysis of interactions between multiple, dynamic organelles over time, both in live cells and by EM.

To demonstrate proof-of-principle for this application, we traced 3 interacting lysosomes over a period of 162s (Figure 5), using the same experimental set up as in the previous experiments (lysosomes stained with SirLyso, ER with mEmerald-Sec61β). The larger size lysosome#1 and relatively smaller lysosome#2 meet at 12s and traffic a short range together until 65s. At 72s, lysosome#3 becomes visible, and all 3 organelles move through separate tracks between 94s-162s (Figure 5a). They all stop moving between 162-180s (Figure 5b, see also Supplementary live-cell Video 8). The movement of lysosome#1 is short-ranged and diffusive, whereas the movements of lysosome#2 and #3 are long-ranged and directional, indicating microtubule-based transport. Whether these lysosomes just interact or partially fuse during the course of live-imaging, is not possible to distinguish by FM. Lysosome#1 and lysosome#2 lack endocytic fiducials, whereas lysosome#3 does display a fluorescent signal from the fiducial particles (Figure 5b). After EM preparation, we used the position information of the fiducials to trace an extremely small volume across the cell (Figure 5c-e) in which we targeted 3 ROIs, each containing 1 identified lysosome, one by one. Each ROI is approximately 2×2 µm (x,y) in size (1.55×1.86×0.98 µm, 1.73×1.68×2.98 µm and 2.16×2.24×1.24 µm exactly), and it required a little over 1 hour (64, 68 and 96 minutes) to collect the complete FIB.SEM datasets. So, within approximately 4 hours we collected 3 distinct FIB.SEM datasets around 3 targeted lysosomes that by live-cell imaging were seen to interact.

**Figure 5.**
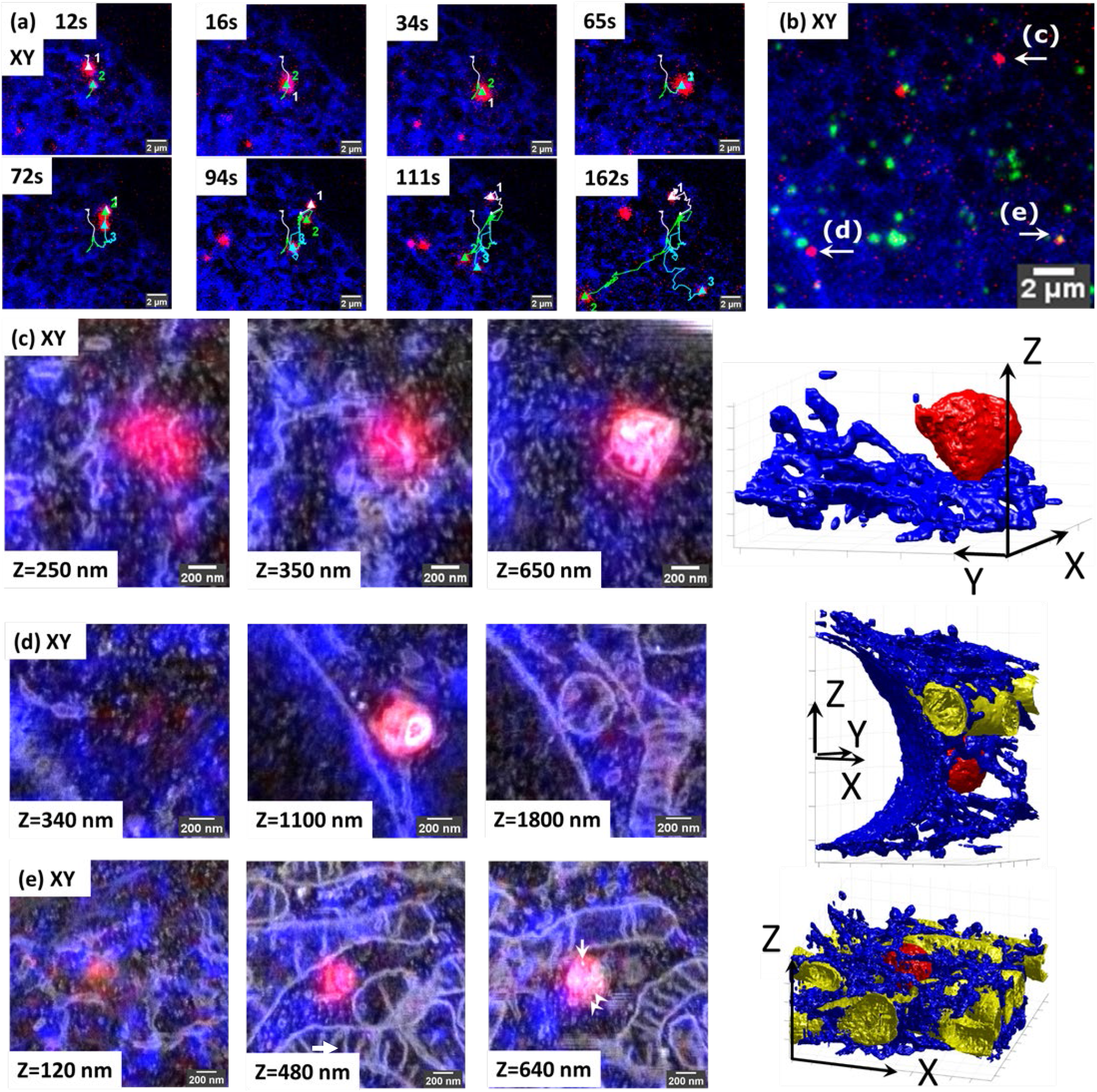
Retracement of interacting organelles by targeted volume-CLEM. (a) Stills from live cell movie of 3 interacting lysosomes showing live lysosomes (SirLyso, red) and ER (mEmerald-Sec61β, blue). The lysosomes are initially visible as separate spots (65s), then split into 2 (72s) and then 3 (94s) spots that travel to different directions. (b) Maximum intensity projection of the Z-stack made after in-situ fixation at 180s. The Z-stack includes endocytosed fiducial particles (green). The 3 interacting lysosomes tracked in live-cell imaging are indicated as c, d and e. (c – e) Each lysosome is traced back for volume-EM, following the targeted imaging routine. Volume-CLEM images of lysosome#1 (c), lysosome#2 (d), lysosome #3 (e), respectively, in 2 xy planes, showing the segmentation of the lysosomes (red), ER (blue), and mitochondria (yellow). Arrow points to the single endocytic fiducial particle, and arrowheads depict the ILVs.

Upon ultrastructural examination and segmentation of the FIB.SEM data, lysosome#1 (red) was found to have a size of 0.56×0.56×0.51 µm x, y, z, with an asymmetrical shape, and positioned at the rim of the cell between the ER (blue) and the plasma membrane. It had little interactions with ER (blue) (Figure 5c, see also Supplementary 3D-overlay Video 9 and Supplementary 3D-segmentation Video 10 and Video 11). Lysosome#2 (red) had a diameter of 0.55×0.58×0.62 µm, with a rather spherical shape, positioned in the perinuclear region, touching the nuclear envelope and making contact with a mitochondrion (yellow) and occasional ER tubules (blue). Its luminal content with degraded membranes was comparable to lysosome#1 (Figure 5d, see also Supplementary 3D-overlay Video 12 and Supplementary 3D-segmentation Video 13 and Video 14 and Video 15). Lastly, lysosome#3 (red) had a diameter of 0.49×0.53×0.46 µm x, y, z, with an almost spherical shape, and positioned in a mitochondria and ER-rich region so dense that the 3D representation became difficult (Figure 5e, see also Supplementary 3D-overlay Video 16 and Supplementary 3D-segmentation Video 17 and Video 18 and Video 19). Lysosome#3 was completely surrounded by mitochondria, which may explain its block in movement between 162-180s. In-contrast to lysosomes #1 and #2, lysosome #3 contains less electron dense material, multiple ILVs (arrowheads) and a single endocytic fiducial particle (arrow, bright spot), defining it as late endosome rather than lysosome.

These data show that targeted volume-CLEM provides a fast and feasible method to study time-resolved interactions between multiple organelles and with high resolution. When combined with overexpression or depletion of specific cargo or transport machinery proteins, this approach will provide a unique way to link molecular regulation to transport dynamics and organelle positioning.

## Discussion

Here, we have established a volume-CLEM pipeline to correlate each and every organelle imaged in live cells to volume-EM with high-throughput and high-precision. We reached this by bringing together recent developments in integrated CLEM instrumentation, using an integrated CLSM-FIB.SEM, novel endocytic fiducial markers that remain fluorescent in resin embedded samples (Fokkema et al., 2018), and advances in sample preparation that minimize resin embedding (Schieber et al., 2017; van Donselaar et al., 2018). Using 3D z-stacks of endocytic fiducial particles as a 3D coordinate anchor between stand-alone CLSM and integrated CLSM platforms, we match the coordinate planes between 3D-FM and 3D-EM modalities, and identify exact coordinates for each and every organelle imaged in live-cell FM back in volume-EM, by FIB.SEM. In **Figure 2**, we show the direct correlation of live-cell imaging data of hydrolase active lysosomes and ER to volume-EM data with a correlation precision far below the x, y, z resolution limits of the FM.

We succeeded to achieve the <100nm sub-organelle level correlation accuracy between 3D-FM and 3D-EM datasets by making use of only a few endocytosed fiducial particles, which function as anchor points between 3D FM and 3D EM. Importantly, correlation is not limited to organelles bearing fiducial particles. Also other fluorescence signals, which are lost after EM embedding, can with high precision be correlated to the volume-EM data, using the fiducials as landmarks. Hence, we are not hindered in our EM analysis by the presence of abundant quantities of fiducial marker. To realize this we made use of Au core-silica shell particles recently developed in our labs which, preserve their fluorescence in epoxy resin (Fokkema et al., 2018). Other fiducial particles with the same properties (Han et al., 2019; Kukulski et al., 2012; Takizawa et al., 2015) are equally suited for our approach. Moreover, other fluorescent probes (Fu et al., 2020; Hemelaar et al., 2017; Morrison et al., 2015; Paez-Segala et al., 2015; Tanida et al., 2020), sample preparation routines (Andrian et al., 2020; Brama et al., 2015; Hohn et al., 2015; C.J. Peddie et al., 2014), and resins (Zhou et al., 2017) developed to retain the available fluorescence signal after EM sample preparation can be adapted and used within the here described pipeline, using fiducials as correlation anchor.

Precise correlation with *a priori* information from live-cell and confocal FM provides exact 3D coordinates of structures within the complete block and enables targeting for volume-EM. In **Figure 3**, we show targeted volume-CLEM (i.e. using the FM identified coordinates to target the ROI in EM) of a single lysosome within the whole cell volume. Blind assessment of the 3D correlation accuracy showed that we can target a single point with sub-100 nm accuracy in x, y, and ∼300 nm accuracy in Z direction. Hence, an ROI identified in CLSM can be correlated, targeted and imaged in volume-EM with at least 100nm confidence. This enables 3D EM visualization of a single organelle, e.g. the lysosome tracked in Figure 3, in a notably short time (∼2.5 hours) and directly correlating to the corresponding FM data. Also addressing homotypic organelle interactions, we investigated prolonged interactions of 3 lysosomes in live cells in **Figure 5**. Subsequent FIB.SEM of each 3 lysosomes one by one was completed in less than 4 hours, highlighting the novel means that targeted volume-CLEM pipeline offers to analyze the ultrastructure of previously interacted organelles in a high throughput manner.

High-throughput volume-CLEM including time-resolved functional imaging in live cells, opens up novel possibilities to study the regulation of rare, transient cellular processes with ultrastructural resolution. Presented as one of our examples, an extensive understanding of MCSs have been uncovered in the last years. In **Figure 4** we provide the first direct evidence that ER-lysosome contacts regulate lysosome movement and shape, in 3D with high temporal and spatial resolution. We analysed the motility of a single lysosome with respect to its interactions with ER in live-cells, and visualized that the lysosome movements mainly follow the surrounding ER tubules. Interaction with dense (bright) ER on one (right) side seemed to obstruct its movement. Once tracked and imaged in volume-EM, we showed that the non-obstructive ER at the left-side ER showed a tubular thin lumen, which only touches the lysosome. In contrast, dense ER on the obstacle-side exhibits a sheet-like thicker lumen and wraps the lysosome surface. Interestingly, the curvature of the lysosome on the ER wrapped site was much flatter compared to its left-site (Figure 4d), indicating a role of ER contacts in the shape and possibly composition of lysosome membrane domains. The data also confirmed the presence of multiple types of contact sites between ER and lysosomes, possibly with different functions (Bonifacino & Neefjes, 2017; Fermie et al., 2018; Garrity et al., 2016; Lim et al., 2019). Numerous proteins involved in MCSs are currently being identified, and many studies focus on addressing the role of these proteins in spatial and temporal regulation of MCSs at the molecular level (Cabukusta et al., 2020; Huang et al., 2020; Lim et al., 2019). We believe our method, uniquely linking live-cell imaging of single organelles to ultrastructural detail, is promising to greatly accelerate understanding in temporal and structural regulation of MCSs at the system and molecular level.

Besides organelle biology, other fields of cell and developmental biology, model organism studies, and clinical studies can also benefit from our novel, targeted volume-CLEM pipeline. A specific development stage in a model organism, a specific cell within an organoid, and a certain cell cycle stage can be selected by live-cell imaging to address the ultrastructural changes in e.g. cellular differentiation, mechanisms of cellular polarization, and cytokinesis at nanometer scale. High precision correlation of 3D-FM and 3D-EM data will allow the use of 3D culture models (e.g. spheroids, organoids) in high-throughput volume-CLEM, which is currently far from trivial (Ando et al., 2018; Rios & Clevers, 2018).

Any fluorescence/light microscopy technique can be incorporated prior to EM sample preparation in the presented volume-CLEM pipeline. We have presented live-cell imaging and confocal FM, which forms the foundation for other FM techniques. Specialized fluorescence methods to study organelle dynamics and membrane trafficking (FRAP), transient molecular interactions (FRET), and local exponential fluorescence decay rates in a sample (FLIM) can be directly incorporated within the workflow (De Los Santos et al., 2015) and also super-resolution techniques to study subcellular structures with greater temporal and/or spatial resolution (Hell et al., 2015; Schermelleh et al., 2019). The registration accuracy is currently limited by the FM resolution, and the volume-CLEM pipeline would clearly benefit from the improved lateral and axial resolution, super-resolution FM could provide (Fu et al., 2020; Shtengel et al., 2014). Similarly, the fluorescence labelling strategies can be adapted to include any types of fluorescent probes, functional reporters (as shown by SirLyso), fluorescent proteins (as shown by mEmerald), and inorganic dyes (as shown by Rhodamine in fiducial particles). The targeted volume-CLEM pipeline is therefore fully flexible in terms of light microscopy approaches.

The targeted imaging strategy reported here would be also very beneficial to unite with another exciting EM technique, cryo electron tomography (cryo-ET) to aid cryo-CLEM workflows. These workflows include freezing the cells or tissues, first imaging them in cryo-FM to identify ROI, then transferring them to cryo-FIB.SEM to prepare lamella from the selected ROI (selected Z-plane), and then finally transferring them to the cryo-TEM to acquire the series of 2D images creating the 3D model. Even though being enormously powerful for studying cellular and molecular interactions in situ with unmatchable resolution, this method is very laborious, low throughput, and available to a limited number of groups in the world. The targeted imaging approach would also solve the main bottleneck also present in these approaches by increasing accuracy and throughput. Implementing a similar iCLSM in a cryo-FIB.SEM set-up would assure the lamella preparation of the correct plane, bearing the ROI, for follow-up cryo-TEM imaging. This would unlock the true potential of cryo-CLEM to identify and study rare cellular processes in-situ, with molecular resolution. Such integrated platforms are currently being commercialized (e.g. METEOR, Delmic).

Structure - function studies of (sub)cellular events require quantitative analyses, which necessitate repetitive imaging of smaller volumes from several independent samples. The here presented targeted volume-EM imaging is a powerful way to find back a small ROI within a large sample. This is especially relevant in a disruptive technique such as FIB.SEM, and very important to unveil the true quantitative potential of volume-EM, which we are just starting to fully utilize (Hoffman et al., 2020; Lucas et al., 2012; Christopher J Peddie & Collinson, 2014). Another related asset of the targeted imaging approach presented here is the time and resource savings it provides. Usually imaging a full mammalian cell with 5nm isotropic resolution in FIB.SEM takes 5-7 days, whereas the majority of the cell volume does not contain relevant information on the dedicated research question. In contrast, our targeted volume-CLEM pipeline can identify, target and image a ROI guaranteed to address the research question within 1-2 hours, with high confidence. The reduced EM imaging duration saves both personnel and machine time, which is crucial for instruments like FIB.SEMs often shared by multiple users in imaging facilities. It also considerably cuts back on the post-collection computational requirements of alignment, reconstruction, segmentation of large 3D-EM data sets (e.g. of whole cells), and the possible correlation with the FM data, which can take as long as the experiment itself. The time for targeting, imaging and correlation could be further shortened by integrating iCLSM and EM operating softwares (e.g. SBEMimage (Titze et al., 2018), MAPS/Thermo Fischer, Atlas/Zeiss, ODEMIS/Delmic). In addition, precise 3D-FM to 3D-EM correlation can facilitate automated image processing, and provide ground-truth for ongoing efforts on automatized segmentation of 3D-EM datasets (Xiao et al., 2018). Together, these will enable fast and user-independent quantification (e.g. volume, membrane interactions) of structures of interest in significant sample sizes.

## Methods

### Cell culture

HeLa cells were cultured in a 37 °C, 5% CO_2_ incubator, in T75 culture bottles (Corning). Cells were maintained in Dulbecco’s Modified Eagle’s Medium (DMEM; Gibco) supplemented with 10% fetal bovine serum, 2 mM L-glutamin, 100 U/mL penicillin, 100 *μ*g/mL streptomycin (referred to as complete DMEM). Cells were passaged when confluency reached 85% to 90%.

For CLEM, Hela cells were grown on gridded glass coverslips, prepared as described earlier (Fermie et al., 2018). On the next day following seeding, the cells were transiently transfected with a construct encoding mEmerald-Sec61b-C1 (Nixon-Abell et al., 2016), which was a gift from Jennifer Lippincott-Schwartz (Addgene plasmid # 90992), for 16 hours. Transfections were performed using Effectene transfection reagent (Qiagen) according to manufacturer’s instructions. Prior to FM, cells were incubated with fiducial markers (Fokkema et al., 2018) at a concentration of 1 *μ*g/ml, and cell permeable lysosome stain SiR-Lysosome (SpiroChrome) at a concentration of 0.5 *μ*M in complete DMEM and incubated for 3 hours.

### Fluorescence Microscopy

Live imaging was performed on a Deltavision RT widefield microscope (GE Healthcare) equipped with a conditioned imaging chamber set to 37°C and 5% CO_2_. Time-lapse imaging was performed using a 100×/1.4 numerical aperture (NA) oil immersion objective and images were recorded on a Cascade II EM-CCD camera (Photometrics). Live-cell imaging was performed for 3-5 minutes, after which the cells were fixed in situ by addition of 1 mL of fixative containing 4% paraformaldehyde (Sigma) and 0.05% glutaraldehyde (25% solution in dH2O, Merck) in 1× PHEM buffer (60 mM PIPES, 25 mM HEPES, 10 mM EGTA, 2 mM MgCl2, pH=6.9) to the imaging holder with the camera still active, to obtain images until the cells are fixed. After fixation, a Z -stack was recorded for all fluorophores using a Zeiss LSM700 CLSM equipped with 63x/1.4 NA oil immersion objective. Z-stacks were collected with 200 nm step size. The position of cells relative to the grid of the coverslips was recorded using polarized light.

### Sample preparation for Volume-EM

Cells were prepared for electron microscopy according to a protocol described earlier, with minor modifications (van Donselaar et al., 2018). Briefly, samples were postfixed using 1% osmium tetroxide (w/v) with 1.5% potassium ferrocyanide (w/v) for 1 h on ice, incubated with 1% thiocarbohydrazide in dH_2_O (w/v) for 15 min, followed by 1% osmium tetroxide in dH_2_O for 30 min. Samples were EM stained with 2% uranyl acetate in dH_2_O for 30 minutes and stained with Walton’s lead aspartate for 30 min at 60 °C. Dehydration was performed using a graded ethanol series. Samples were embedded in Spurr resin and polymerized for 48–60 h at 65 °C following the extremely thin layer plastification method(van Donselaar et al., 2018). Resin embedded samples on the glass coverslips were subsequently coated with 8 nm carbon and carbon-tape mounted on aluminum stubs.

### Electron Microscopy

A Scios FIB.SEM (ThermoFisher) was used. It included an Everhard-Thorley Detector (ETD), an in-lens detector of Back Scattered Electrons (BSE), and an in-the-column detector of Secondary Electrons (SE). The ETD was used for imaging of sample surfaces. The BSE detector was used for the imaging cellular ultrastructure in 3D. The in-the-column SE detector was used to enhance fiducial marker contrast. The FIB was equipped with a Ga-ion source; for the 3D acquisition a current of 0.4 nA and an acceleration voltage of 30 kV were used. The 3D acquisition in the FIB.SEM was controlled by the Slice&View software version 3 (ThermomFischer).

### Integrated CLSM

The integrated CLSM was equipped with a Nikon industrial inspection objective lens (ELWD series, Plan Apo 100x NA0.9). A lens of focal length 120 mm (25 mm diameter, OptoSigma anti-reflection coated doublet lens DLB-25-120PM) was used as a tube lens. For lateral optical scanning a Yanus scan-head (FEI Munich) equipped with a 50 mm focal length scan-lens was used. As an excitation source a 532 nm laser (Omicron, integrated with an Acoustic Optical Modulator in LightHUB housing) coupled via a single-mode optical fiber was used. The excitation and detection light paths were combined inside the scanning head via a dichroic mirror (T560lpxr-UF2, Chroma). The detected light was coupled into a multi-mode fiber (FG010LDA, Thorlabs) with a core diameter of 10 µm (about 1 Airy disk). The light from the sample was split by a second dichroic mirror (T565LP, Chroma) into a fluorescence and a reflection parts. Back-reflected light was detected by a PMT operating in current mode (PMMA01, Thorlabs). The fluorescence signal was, after bandpass filtering (ET585pxr-65, Chroma), detected by a photon counting PMT (H7422P-40, Hamamatsu). For sub-micrometer axial scanning a single axis piezo-stage (E-601.1SL, Physik Instrumente) was mounted on top of the FIB.SEM stage. iCLSM was controlled via a National Instruments NI-6251 DAQ card using LabView software. See also Supplementary Figure 1 for the iCLSM/ FIB.SEM set-up.

### Integrated CLSM and FIB.SEM Pipeline

Initially a low magnification, low exposure SEM image was acquired in order to find the main grid markings (numbers and letters). Using those markings the square where the life cell imaging was performed was found. Another low exposure SEM image (“snapshot” at current of 10 pA, dwell time 0.5 us and pixel size of ∼250×250 nm^2^) was taken of the suspected square in order to verify the correct cells were found. Performing of these steps using SEM has advantage because of the much wider field of view of SEM compared to that of the integrated CLSM (up to 350 um), and of the large depth-of-focus of the SEM. However, the topographical contrast of the SEM is much lower than of the reflection light channel of the integrated CLSM. The SEM exposure of the sample should be minimized as much as possible since the fluorescence quenches strongly under electron radiation. On the next step the sample was transferred under the integrated CLSM. Here we made an overview images with pixel size larger than 0.5 µm and dwell time of 3 µs. In the reflected light channel we can find the cells and refine the Z position of the sample. The shape of the cell is used then to find ROI from the life-cell/fixed cell images. Next we take a fine resolution image (34×34 nm^2^ pixels, Z step of 100 nm and dwell time of 24 µs) of cell area of 35×35 µm^2^ or smaller. In the fluorescence channel in the acquired image some of the nano-fiducial particles were visible. Their configuration allowed to refine the position of the ROI relative to the surface features detected with the reflection light channel. All ROIs selected in the life cell data were recorded in the same manner with the integrated CLSM. Next, the sample was transferred back under the FIB.SEM. At this step the surface features of the cells were accurately studied in SEM with pixel size of 17×17 nm and dwell time of 1 us and beam current of 50 pA, since fluorescence quenching had become of no issue after the integrated CLSM imaging was performed. The surface features of the cell recorded at this step were used in order to refine the ROI in the SEM field of view and to determine the acquisition volumes of the FIB/SEM itself.

### Alignment of FIB.SEM data

The raw SEM images of 3D FIB.SEM data were aligned using MatLab. The vertical alignment (Y axis in SEM images, Z axis in the original orientation) was performed via the detection of the glass substrate in the SEM images. The alignment in the X direction was performed using correlation of consecutive images. The height of XY slices is calculated relative to the substrate level.

### Correlation/ Targeted Imaging Routine

In the live-cell data the ER and the lysosomes are used to define the region of interest. After fixation, CLSM image is collected of the same lysosomes and the ER, and additionally the fiducial particles emitting in another channel. The shape of the cell visible in SEM and the ER configuration can be used for correlation. The fluorescence of the nano-particles is visible in the integrated CLSM and allows for registration with the non-integrated CLSM. The fluorescence channel of integrated CLSM is directly linked with the reflection channel of the integrated CLSM. In the reflected light channel the cell shape and cell topography is visible in fine detail. The same fine details (i.e. slopes, protrusions and bumps) are visible in the reflection light channel of the integrated CLSM and in the SEM image. This fine details can be used for the fine registration between these two (vacuum) modalities. Thus the accurate finding of the ROI in the FIB.SEM relies on the following ladder of registration steps: from the fiducials in the non-integrated CLSM to the fiducials visible in the integrated CLSM. Next, the fiducials in the CLSM can be related exactly to the surface features of the cell, with are also visible in the SEM albeit with a different contrast.

### Image Correlation, Analysis, and Segmentation

Registration of 3D CLSM and 3D FIB.SEM data was achieved by the following steps: First, 2D maximum intensity projection of CLSM data and 2D maximum intensity projection of integrated CLSM data were registered manually to find relative rotation angle. The registration was based on the clusters of fluorescent nanoparticles. Next the whole 3D CLSM data was rotated around Z axis by the angle found at the first step. Then the sub-volume of 3D CLSM enclosing the ROI was copied with a several pixel margin. Afterwards, the sub-volume was interpolated to a smaller voxel size using bilinear algorithm. The 3D FIB.SEM data were reordered to match those of CLSM data (the native orientation in Slice&View software is XZY) and voxel dimensions were equalized: binned along X and Z directions (SEM scanning directions) and interpolated along Y direction (FIB slicing direction). The voxel sizes were matching those of 3D CLSM data sub-volume at the previous step. Finally, the correct sub-volume of 3D CLSM data was selected manually, and the 3D FIB.SEM data was added to it as a 4^th^ color channel.

We employed a semi-automated approach of the custom written MatLab code (see Supplementary information, and Supplementary Figure 3) for segmentation of the 3D FIB/SEM datasets. The segmentation was performed consequently for each type of cell compartments. At first, the lysosomes were segmented. Then the voxels of the lysosome were zeroed in the volume and the segmentation was performed for the mitochondria (if any were present in the volume). As the last step the ER segmentation was performed with all the previously segmented organelles zeroed in the volume.

The semi-automated segmentation “in bunches” was performed in the following manner. For every 10 2D slices the average image (i.e. “bunched slices”) was shown to the user. Then the user drew a contour around the suspected organelle which defined the area where the automated segmentation would be performed. All automated steps were performed on 2D basis using the standard MatLab Image Processing Toolbox functions. Otsu thresholding was performed (2 levels in case of lysosome and 3 levels in other cases) in the user selected area of the averaged image. The highest level obtained from the Otsu algorithm-was used for thresholding the suspect area of each of the substitute original slices of the stack. The images were then subjected to Canny edge detection, and the edges were morphologically filled. The automated steps were parallelized for the slices of the bunch. After the automated steps were performed, the new averaged image was shown to the user with the detected areas zeroed. At this step the user could define another area where the organelle was visible or accept the results of the automated steps and proceed to process the next bunch of the slices.

For each organelle the segmentation procedure was performed three times along three main axis the stack (for XY, XZ and ZY slices). A pixel was considered belonging to the organelle if it was detected in any of the two steps.

## Supporting information

Supplementary 3D-overlay Video 1

Supplementary live-cell Video 2

Supplementary 3D-overlay Video 3

Supplementary live-cell Video 4

Supplementary 3D-overlay Video 5

Supplementary 3D-segmentation Video 6

Supplementary 3D-segmentation Video 7

Supplementary live-cell Video 8

Supplementary 3D-overlay Video 9

Supplementary 3D-segmentation Video 10

Supplementary 3D-segmentation Video 11

Supplementary 3D-overlay Video 12

Supplementary 3D-segmentation Video 13

Supplementary 3D-segmentation Video 14

Supplementary 3D-segmentation Video 15

Supplementary 3D-overlay Video 16

Supplementary 3D-segmentation Video 17

Supplementary 3D-segmentation Video 18

Supplementary 3D-segmentation Video 19

## Acknowledgments

This work was supported by funding from Stichting voor de Technische Wetenschappen (STW), grant number 12715, granted to HG and JK. NL also acknowledges the Netherlands Organization for Scientific Research (NWO) for an ZonMW-TOP grant awarded to JK. Authors also acknowledge NWO roadmap grant The Netherlands Electron Microscopy Infrastructure (NEMI) chaired by JK.

## Author Contributions

NL, SL, JFe, HG, and JK designed the study. JFe performed the cell culture, labelling, live-cell imaging, confocal microscopy, and analysed the data with NL. JFo synthesized, characterized and provided the endocytic fiducials. JFe, CdeH optimized and performed EM sample preparation. SL, AA, GB, and HG designed and built the integrated CLSM. SL performed integrated CLSM and FIB.SEM imaging, and analysed the data with NL. SL did the 3D image correlation and segmentation, and prepared the figures with NL. NL and SG wrote the paper with input from all authors. HG and JK reviewed the manuscript.

