## Supplementary Information for "Correlative Organelle Microscopy: fluorescence guided volume electron microscopy of intracellular processes"

### List of Supplementary Videos

1. Supplementary 3D-overlay Video 1 (related to Figure 1i &j and Figure 2d)
2. Supplementary live-cell Video 2 (related to Figure 2a)
3. Supplementary 3D-overlay Video 3 (related to Figure 3d)
4. Supplementary live-cell Video 4 (related to Figure 4a)
5. Supplementary 3D-overlay Video 5 (related to Figure 4d):
6. Supplementary 3D-segmentation Video 6 (related to Figure 4e)
7. Supplementary 3D-segmentation Video 7 (related to Figure 4e)
8. Supplementary live-cell Video 8 (related to Figure 5a and 5b)
9. Supplementary 3D-overlay Video 9 (related to Figure 5c)
10. Supplementary 3D-segmentation Video 10 (related to Figure 5c)
11. Supplementary 3D-segmentation Video 11 (related to Figure 5c)
12. Supplementary 3D-overlay Video 12 (related to Figure 5d)
13. Supplementary 3D-segmentation Video 13 (related to Figure 5d)
14. Supplementary 3D-segmentation Video 14(related to Figure 5d)
15. Supplementary 3D-segmentation Video 15 (related to Figure 5d)
16. Supplementary 3D-overlay Video 16 (related to Figure 5e)
17. Supplementary 3D-segmentation Video 17 (related to Figure 5e)
18. Supplementary 3D-segmentation Video 18 (related to Figure 5e)
19. Supplementary 3D-segmentation Video 19 (related to Figure 5e)

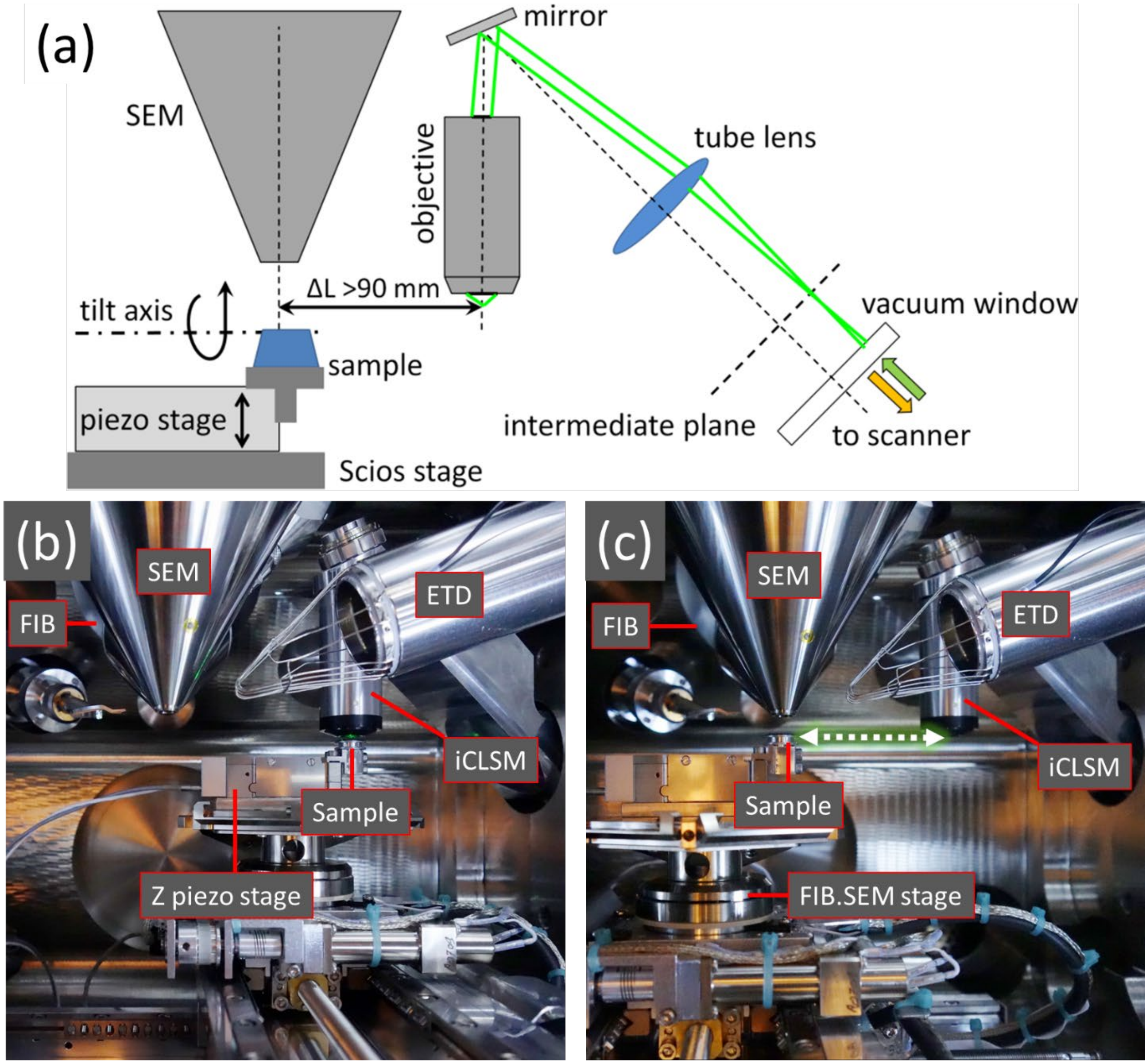


**Supplementary Figure 1. The integrated CLSM- FIB.SEM setup** built in a FEI Scios system as seen from the right wall of the vacuum chamber. **(a)** iCLSM and FIB.SEM observe the sample from the same direction Samples can be investigated with the integrated CLSM by bringing the sample under the objective lens of the iCLSM. **(b)** Switching between iCLSM and (FIB.)SEM imaging is accomplished by stage translation using an accurate motorised stage of the FIB.SEM system.

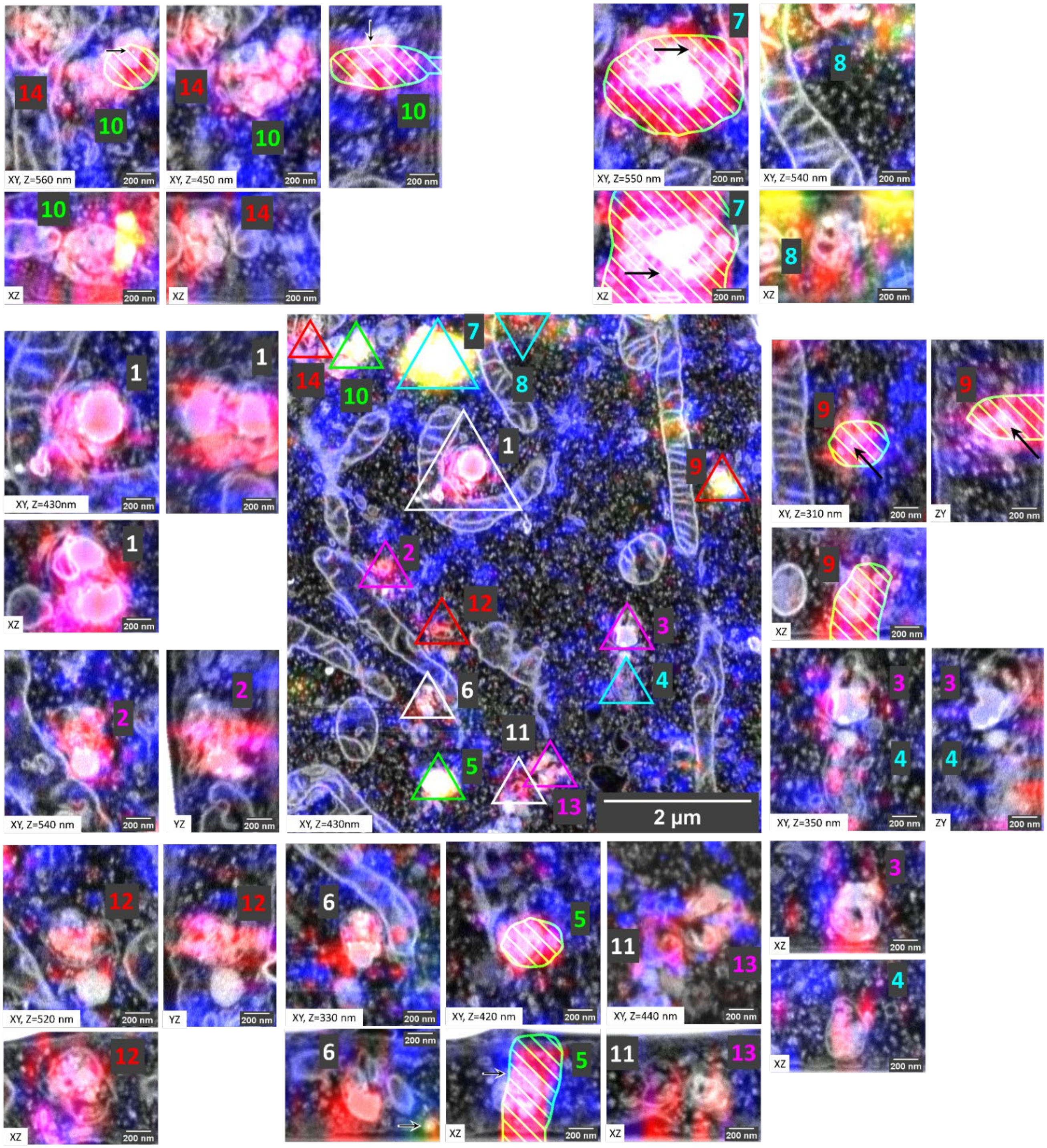


**Supplementary Figure 2.** Fluorescence data of every single live-cell imaged organelle in the ROI can be correlated to ultrastructural 3D-EM data with high precision. The cells have endocytosed fiducial particles (green), mEmerald-Sec61β is localized to ER (blue), and lysosomes are stained with SirLyso (red). Enzymatic activity of 14 lysosomes has been correlated to their ultrastructure.

**Supplementary Information on Segmentation and Visualization**

The segmentation program is implemented to segment a single organelle type in one run. If some of organelles were already segmented those voxels are suppressed in the present run (see the dark region in **Supplementary Figure 3**, corresponding to the previously segmented lysosome). The direction of slicing can be selected as XY or YZ or XZ slices. The best segmentation results are achieved when the segmentation is performed along all 3 possible directions. The results of these three runs is merged: a voxel is said to belong to the segmented organelle if it appears in any two of the three segmentation runs.

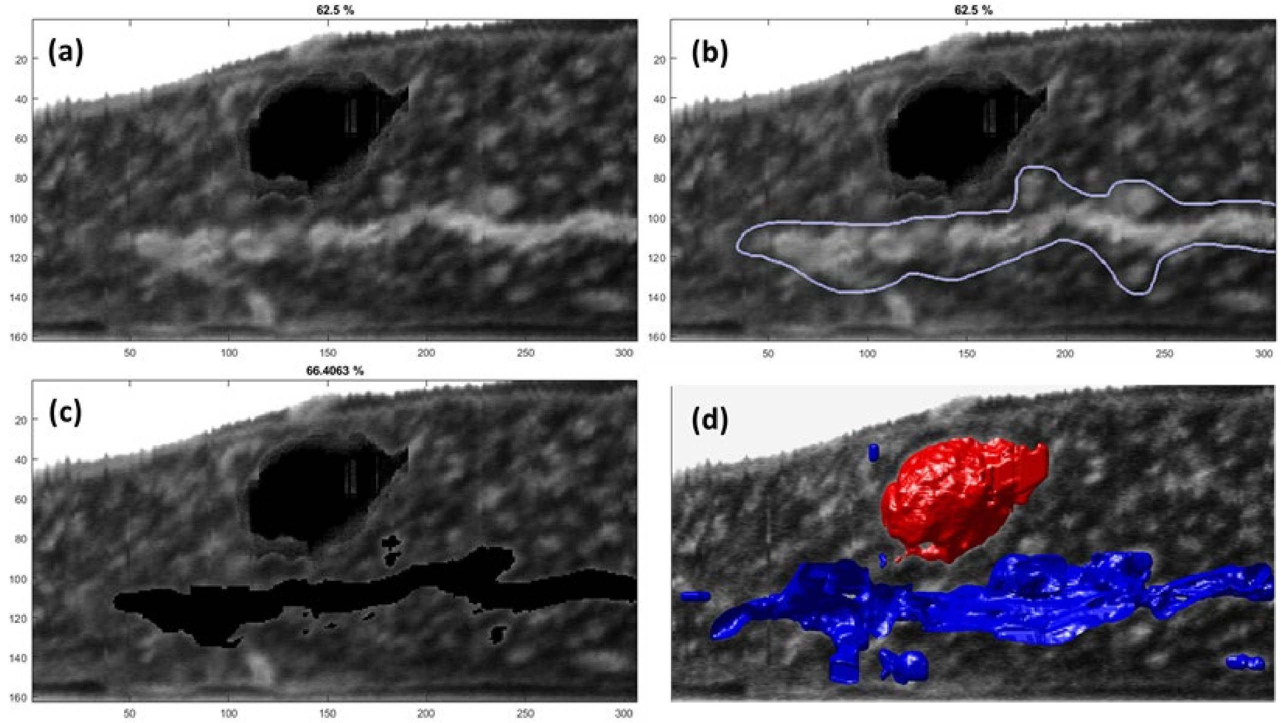


**Supplementary Figure 3. (a)** The average intensity of 10 XZ slices of a 3D Stack are presented to the user. The dark area at the top part of the cell is a lysosome segmented at the previous segmentation stage. **(b)** The area where the ER is predicted is selected by the user and highlighted with blue line. **(c)** Automatic segmentation was performed over the manually selected area on all of the slices of the image bunch. The pixels selected are set to zero. The user can now define a new area for segmentation on the same bunch or process to the next bunch of slices. **(d)** Segmented organelles are visualized in 3D.

The user is presented with averaged 10 consecutive 2D slices (i.e. bunched slices) in a full 3D stack. The manually selected area where the organelle of interest is visible is then subjected to automated segmentation. Each of the slices of the bunch is processed automatically by applying intensity threshold and edge detection. Next, the image of the same bunch (excluding just segmented areas) is shown again to the user to select another piece of the organelle. When there are no more non-segmented areas remain in the current bunch of slices, the user can double click) to proceed to the next bunch.

The program was implemented in MatLab with Image Processing Toolbox and Parallel Computing Toolbox. Find the code below:

```
bunching=10;% how many slices are bunched together
order=[3,1,2];% dimensions during processing, [3,1,2] means XZ slices
Stack=permute(Stack_0.*uint8(~(Stack_V_OR)),order);%3D Stack with previously segmented parts
excluded
Stack_segmented=zeros(size(Stack),'logical'); % empty segmentation variable
N=size(Stack,3);% total number of 2D slices
start=1;% in case some first slices are known to be empty
close all; figure;
for i=(1:(ceil((N-start)/bunching)))% main cycle
    I_bunch=Stack(:,:,(start+(i-1)*bunching):min(start+i*bunching,N));% current bunch of
slices
    I_0=mean(I_bunch,3);%image to show - average intensity over the bunch

    imagesc(I_0);axis tight;colormap gray;% show new image in black-and-white
    title([num2str(i*bunching/N*100),' %']);% show progress in percent

    mask_bunch=zeros(size(I_bunch),'logical');%variables for bunch segmentation
    I_detected=mask_bunch;I_t=mask_bunch;

    size_check=1; % variable to track
    while size_check %% process membrane pieces one-by-one in the current bunch
    ans=[];
        while isempty(ans) %% this syntax allows to redraw manual selection
            h = imfreehand();% user highlights area
            wait(h);
        end
I_manual=createMask(h); % the area selected by user
        if (sum(double(I_manual(:)))>10) %selected area is large

          treshold=multithresh(I_0.*double(I_manual),2);%for endosome - 1, for ER - 2
          parfor j=1:bunching % automated segmentation for individual slices
            I_t=I_bunch(:,:,j)...% one slice of the bunch
                .*uint8((I_bunch(:,:,j)>treshold(numel(treshold))).*I_manual);%tresholded
            I_detected= imerode(imfill( edge( I_t,'Canny',0.7,5)+(I_t>0),'holes'),ones(3));%
            mask_bunch(:,:,j)=mask_bunch(:,:,j)+I_detected;
          end
         %show bunch projection again without segmented area:
         imagesc(double(~max(mask_bunch,[],3)).*I_0);
         axis tight;
         title([num2str((i+1)*bunching/N*100),' %']);%show progress in percent
        else % small area was selected (double click) - go to the next bunch
            size_check=0;
        end
    end
    Stack_segmented(:,:,(start+(i-1)*bunching):(start+i*bunching))=mask_bunch;
end

Stack_segmented=ipermute(Stack_segmented,order);% rearrange dimensions back
```

Example of merging results of 3 segmentation runs:

```
Stack_Mito=(Stack_Mito_123|Stack_Mito_321)&(Stack_Mito_123|Stack_Mito_312)&(Stack_Mito_321|St
ack_Mito_312);
```

For visualization the following commands were used:

```
p = patch(isosurface(double(smooth3(fliplr(Stack_Mito(:,:,295:-1:1))))));
p.FaceColor = 'y';
p.EdgeColor = 'none';
daspect([1 1 27/128]);
axis equal;
camlight;
lighting phong
```
